# The Slit-binding Ig1 domain is required for multiple axon guidance activities of *Drosophila* Robo2

**DOI:** 10.1101/2021.05.07.443153

**Authors:** LaFreda J. Howard, Marie C. Reichert, Timothy A. Evans

## Abstract

*Drosophila* Robo2 is a member of the evolutionarily conserved Roundabout (Robo) family of axon guidance receptors. The canonical role of Robo receptors is to signal midline repulsion in response to their cognate Slit ligands, which bind to the N-terminal Ig1 domain in most Robo family members. In the *Drosophila* embryonic ventral nerve cord, Robo1 and Robo2 cooperate to signal Slit-dependent midline repulsion, while Robo2 also regulates the medial-lateral position of longitudinal axon pathways and acts non-autonomously to promote midline crossing of commissural axons. Although it is clear that Robo2 signals midline repulsion in response to Slit, it is less clear whether Robo2’s other activities are also Slit-dependent. To determine which of Robo2’s axon guidance roles depend on its Slit-binding Ig1 domain, we have used a CRISPR/Cas9-based strategy replace the endogenous *robo2* gene with a *robo2* variant from which the Ig1 domain has been deleted *(robo2ΔIg1)*. We compare the expression and localization of Robo2ΔIg1 protein with that of full-length Robo2 in embryonic neurons *in vivo*, and examine its ability to substitute for Robo2 to mediate midline repulsion and lateral axon pathway formation. We find that removal of the Ig1 domain from Robo2ΔIg1 disrupts both of these axon guidance activities. In addition, we find that the Ig1 domain of Robo2 is required for its proper subcellular localization in embryonic neurons, a role that is not shared by the Ig1 domain of Robo1. Finally, we report that although FasII-positive lateral axons are misguided in embryos expressing Robo2ΔIg1, the axons that normally express Robo2 are correctly guided to the lateral zone, suggesting that Robo2 may guide lateral longitudinal axons through a cell non-autonomous mechanism.

## Introduction

Axon guidance receptors of the Roundabout (Robo) family are widely conserved among bilaterian animals, and their canonical role is to regulate midline crossing of axons by signaling midline repulsion in response to Slit ligands. In groups such as insects and vertebrates, where multiple family members are present, some Robo receptors have acquired additional or alternative activities. In *Drosophila*, three Robo family members (Robo1, Robo2, and Robo3) regulate multiple axon guidance decisions during development of the embryonic ventral nerve cord (VNC). Robo1 and Robo2 cooperate to signal midline repulsion in ipsilateral and post-crossing commissural axons (Rajagopalan *et al*. 2000a; Simpson *et al*. 2000b), Robo2 and Robo3 regulate the medial-lateral position of longitudinal axon tracts (Rajagopalan *et al*. 2000b; Simpson *et al*. 2000a; Spitzweck *et al*. 2010; Evans and Bashaw 2010), and Robo2 promotes midline crossing of commissural axons during the early stages of axon guidance (Simpson *et al*. 2000b; Spitzweck *et al*. 2010; Evans and Bashaw 2010; Evans *et al*. 2015). In some contexts, *Drosophila* Robo receptors (in particular Robo2) can influence development in ways other than by acting as canonical midline repulsive Slit receptors (Kramer *et al*. 2001; Kraut and Zinn 2004; Mellert *et al*. 2009; Evans *et al*. 2015; Ordan and Volk 2015; Alavi *et al*. 2016), but the precise mechanism(s) by which they carry out these additional activities, and whether all of these activities are dependent on interaction with Slit, is not fully understood.

### Structure of Robo receptors and functions of individual receptor domains

Most Robo receptors, including the three *Drosophila* Robos, share a characteristic arrangement of eight extracellular structural domains: five immunoglobulin-like domains (Ig1-Ig5) plus three fibronectin type III domains (Fn1-Fn3). The cytoplasmic regions of Robo receptors are more divergent, but share some or all of four conserved cytoplasmic (CC) amino acid motifs (CC0-CC3). Specific biochemical roles have been identified for some individual ectodomain elements: the N-terminal Ig1 domain is the primary Slit-binding domain in most Robo receptors (Liu *et al*. 2004; Morlot *et al*. 2007; Fukuhara *et al*. 2008; Evans *et al*. 2015; Brown *et al*. 2015), while other domains have been shown to contribute to receptor multimerization (e.g. Ig3 of *Drosophila* Robo2 (Evans and Bashaw 2010) and Ig1, Ig3, and Ig4 of human Robo1 (Aleksandrova *et al*. 2017)) or receptor-receptor interactions (e.g. Ig1-Ig2 of *Drosophila* Robo2, which mediate binding to *Drosophila* Robo1 (Evans *et al*. 2015)). The Ig1 domain of the Robo3/Rig-1 receptor in mammals has lost the ability to bind Slit (Zelina *et al*. 2014), but the receptor has acquired a novel ligand (NELL2) which interacts with one or more of Robo3/Rig-1’s Fn domains (Jaworski *et al*. 2015).

We have previously carried out a comprehensive structure/function study of the ectodomain elements within the *Drosophila* Robo1 receptor, and we found that while the midline repulsive activity of *Drosophila* Robo1 is strictly dependent on its Slit-binding Ig1 domain, each of its other seven ectodomain elements (Ig2-5, Fn1-3) are individually dispensable for midline repulsion (Brown *et al*. 2015; Reichert *et al*. 2016; Brown *et al*. 2018). Although not required for midline repulsive signaling, Robo1’s Fn domains are necessary for its negative regulation by Commissureless (Comm) and Robo2 (Brown *et al*. 2018; Brown and Evans 2020). It is not yet clear precisely which domains in *Drosophila* Robo2 and Robo3 contribute to each of their divergent axon guidance roles, although previous gain of function studies indicate that Robo2’s midline repulsion activity depends on Ig1, its lateral positioning role depends on Ig1 and Ig3, and both Ig1 and Ig2 contribute to its pro-midline crossing activity (Evans and Bashaw 2010; Evans *et al*. 2015).

### Multiple axon guidance roles of *Drosophila* Robo2

Robo2 regulates multiple axon guidance outcomes during development of the *Drosophila* embryonic CNS: 1) it prevents midline crossing of ipsilateral and post-crossing commissural axons in response to the repellant ligand Slit, 2) it promotes midline crossing of commissural axons non-autonomously by antagonizing Slit-Robo1 repulsion, and 3) it regulates the medial-lateral position of longitudinal axon pathways. Robo2 acts alongside Robo1 to signal midline repulsion during the early stages of axon guidance in the embryonic VNC. Genetic data show that this activity of Robo2 is Slit-dependent, as *robo1,robo2* double mutants display more severe midline crossing defects than *robo1* or *robo2* single mutants, and the *robo1,robo2* double mutants phenocopy *slit* null mutants (Rajagopalan *et al*. 2000a; Simpson *et al*. 2000b). However, Robo1 and Robo2 signaling mechanisms are not entirely the same, as *robo1* can rescue *robo2’s* midline repulsive role, but *robo2* cannot substitute for *robo1* in this context (Spitzweck *et al*. 2010).

In addition to its canonical role in midline repulsion, Robo2 also acts non-autonomously to inhibit Slit-Robo1 repulsion in trans to promote midline crossing of commissural axons in the embryonic VNC. We have previously shown that deletion of the Slit-binding Ig1 domain decreases, but does not eliminate, Robo2’s ability to promote midline crossing in gain-of-function experiments (Evans *et al*. 2015).

The mechanism by which Robo2 promotes lateral pathway formation has not been characterized, although it has been proposed that the three *Drosophila* Robo receptors act (either alone or in combination) to specify the medial-lateral distance of longitudinal axon pathways from the midline in response to a midline-secreted Slit gradient (Rajagopalan *et al*. 2000b; Simpson *et al*. 2000a). We have previously shown that Robo2’s ability to induce lateral shifting of medial longitudinal neurons in gain of function experiments is disrupted when Ig1+Ig2 of Robo2 are deleted, consistent with the hypothesis that this activity is Slit-dependent (Evans and Bashaw 2010). If this is the case, we should expect that Slit binding via the Ig1 domain will be required for Robo2’s endogenous lateral positioning activity.

In order to determine the requirements for individual ectodomain elements for the various axon guidance roles of *Drosophila* Robo2, and to distinguish between its Slit-dependent and Slit-independent activities (if any), we have begun a systematic structure/function analysis of ectodomain elements within Robo2. Here, we describe our initial set of experiments using a CRISPR/Cas9-based gene replacement approach to examine the requirement for the Robo2 Ig1 domain for the receptor’s endogenous roles in midline repulsion and lateral pathway formation. We show that each of these activities are disrupted by deletion of the Robo2 Ig1 domain, and we also show that, in contrast to Robo1, the Ig1 domain of Robo2 is also important for proper axonal localization of the Robo2 protein in embryonic neurons in vivo.

## Results

### CRISPR/Cas9-based gene replacement of *robo2*

To begin our functional analysis of the Robo2 ectodomain, we used a CRISPR/Cas9-based gene modification approach (Gratz *et al*. 2014; Port *et al*. 2014) to modify the *robo2* locus to express structural variants of Robo2 (Fig. 1). We first used this approach to create a full-length *robo2*^*robo2*^ allele, in which exons 2-14 of the *robo2* locus are replaced by an HA-tagged full-length *robo2* cDNA. In our *robo2* modified alleles, the endogenous *robo2* promoter, transcriptional start site, first exon (including the start codon and signal sequence), and first intron remain unmodified. Spitzweck et al (2010) used a knock-in approach to similarly replace *robo2* with a full-length *robo2* cDNA, and showed that HA-tagged Robo2 protein expressed from this modified locus was properly expressed and could fully rescue Robo2’s roles in midline repulsion, lateral pathway formation, and promotion of midline crossing (Spitzweck *et al*. 2010).

**Figure 1.**
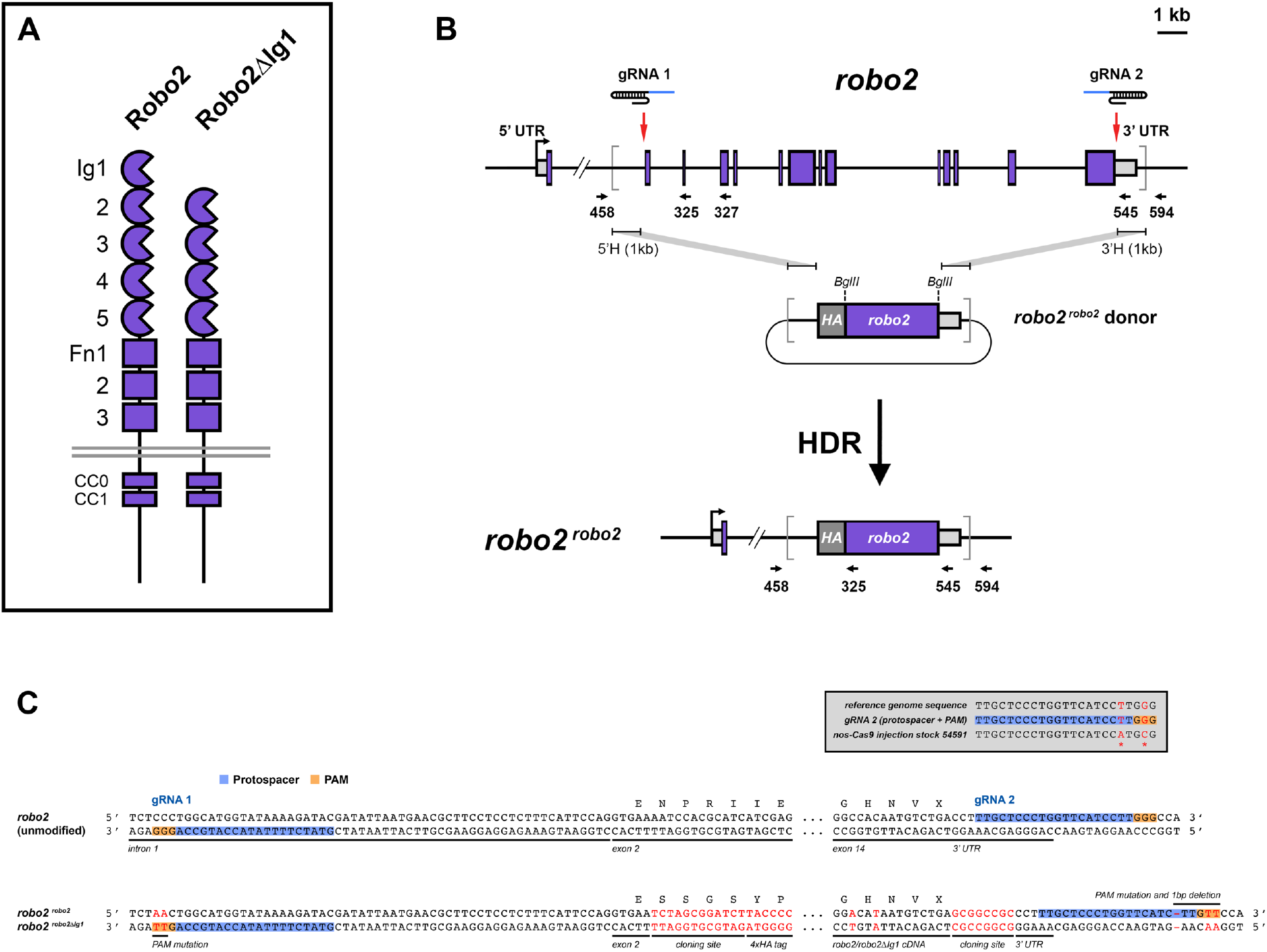
CRISPR/Cas9-based gene replacement of *robo2*. (A) Schematics of the full-length Robo2 protein and the Robo2ΔIg1 variant, from which the Slit-binding Ig1 domain has been deleted. (B) Schematic of the *Drosophila robo2* gene showing intron/exon structure, location of gRNA target sites, *robo2*^*robo2*^ homologous donor plasmid, and the resulting *robo2*^*robo2*^ HDR allele. Endogenous *robo2* coding exons are shown as purple boxes; 5’ and 3’ untranslated regions are shown as light grey boxes. The start of transcription is indicated by the bent arrow. Introns and exons are shown to scale, with the exception of the first intron, from which approximately 19 kb has been omitted. Red arrows indicate the location of upstream (gRNA 1) and downstream (gRNA 2) gRNA target sites. Grey brackets demarcate the region to be replaced by sequences from the donor plasmid. Arrows indicate the position and orientation of PCR primers. The same two gRNAs were combined with a *robo2*^*robo2ΔIg1*^ donor plasmid to create the *robo2*^*robo2ΔIg1*^ HDR allele. (C) Partial DNA sequences of the unmodified *robo2* gene and the modified *robo2*^*robo2*^ *and robo2*^*robo2ΔIg1*^ HDR alleles. Black letters indicated endogenous DNA sequence; red letters indicate exogenous sequence. Both DNA strands are illustrated. The gRNA protospacer and PAM sequences are indicated for both gRNAs. The first five base pairs of *robo2* exon 2 are unaltered in both modified alleles, and the *robo2* coding sequence beginning with codon N90 is replaced by the HA-tagged full-length *robo2* (for *robo2*^*robo2*^) or *robo2ΔIg1* (for *robo2*^*robo2ΔIg1*^) cDNAs. The endogenous *robo2* transcription start site, ATG start codon, and signal peptide are retained unmodified in exon 1. The PAM sequences for both gRNA targets and the protospacer sequence for the gRNA2 target are modified in the donor plasmids, ensuring that the *robo2*^*robo2*^ and *robo2*^*robo2ΔIg1*^ donor plasmids and modified alleles are not cleaved by Cas9. The grey box shows *robo2* sequence polymorphisms present in the *nos-Cas9* injection stock compared to the reference genome sequence, which are predicted to interfere with Cas9 cleavage at the gRNA2 target site. UTR, untranslated regions; 5’H, 5’ homology region; 3’H, 3’ homology region; HA, hemagglutinin epitope tag; gRNA, guide RNA; HDR, homology directed repair; PAM, protospacer adjacent motif.

We generated a guide RNA (gRNA) expression plasmid using the pCFD4 gRNA backbone (Port *et al*. 2014), containing two gRNA sequences targeting the first intron (∼50 bp upstream of exon 2) and exon 14 (3’ UTR) of *robo2*. We also created a *robo2*^*robo2*^ homologous donor plasmid containing the HA-tagged *robo2* coding sequence along with 1 kb upstream (5’H) and downstream (3’H) flanking sequences to serve as a template for homology-directed repair (HDR). The *robo2* coding sequence in this donor construct is flanked by restriction sites, allowing us to swap out the full-length *robo2* sequence for any alternative coding sequence. Using this approach, we should be able to generate many different *robo2* gene replacement variants using the same set of gRNAs and the same homologous donor backbone. For the CRISPR modified alleles described here (*robo2*^*robo2*^ and *robo2*^*robo2ΔIg1*^), each donor construct was co-injected along with the pCFD4 gRNA construct into *Drosophila* embryos expressing Cas9 under the control of the germline-specific *nanos* promoter (*nos-Cas9*.*P*) (Port *et al*. 2014), and F1 progeny from the injected flies were screened by PCR to identify those carrying the expected modification. We generated stable lines from positive F1 flies and sequenced the modified locus fully from at least two lines for each modified allele. Additional details are provided in the Methods.

### *robo2* gRNA target polymorphisms and variable HDR replacements

In addition to the correctly modified *robo2* alleles we recovered for the *robo2*^*robo2*^ and *robo2*^*robo2ΔIg1*^ HDR gene replacements, we also recovered lines that tested positive in our initial PCR screening but deviated from the expected HDR replacements in several ways. These deviations included: variations in the number of N-terminal HA repeats (*robo2*^*robo2*^ line B7-3 had 5xHA instead of 4xHA), deletions within the donor coding sequence (*robo2*^*robo2ΔIg1*^ line T-8 had a 999 bp internal deletion in the Ig4-Fn2 region), and partial replacements that retained all or part of the last exon (*robo2*^*robo2*^ line B7-3 had all introns removed, but the cloning site and modifications to the 3’ end of the cDNA present in the HDR donor were not present in the HDR allele, suggesting that the gene replacement ended somewhere within the final coding exon). Sequencing of genomic DNA fragments from flies in which the 3’ end of the *robo2* gene was not replaced revealed sequence polymorphisms relative to the reference genome sequence for *robo2* (Fig. 1C) which altered the predicted gRNA 2 target site. We infer that these sequence polymorphisms were present in the *nos-Cas9* injection stock and prevented Cas9 cleavage at this site in some or all of the injected flies, which may account for the variations in the extent of the gene replacement at the 3’ end of *robo2*. For the protein expression and phenotypic analyses described below, we used lines in which the replacement was complete and correct, as confirmed by DNA sequencing of the entire modified locus in each line (*robo2*^*robo2*^ line B2-2 and *robo2*^*robo2ΔIg1*^ line O3).

### Expression and localization of Robo2ΔIg1 in embryonic neurons

We have previously shown that deleting the Ig1 domain from *Drosophila* Robo1 does not affect its expression pattern, axonal localization, clearance from commissural axon segments, or regulation by the endosomal sorting protein Commissureless (Comm) (Brown *et al*. 2015). To examine whether the Ig1 domain of Robo2 is similarly dispensable for its expression and localization in embryonic neurons, we used an antibody against the N-terminal HA tag to compare the expression of full-length Robo2 and Robo2ΔIg1 proteins in the ventral nerve cord (VNC) of late-stage *Drosophila* embryos (stage 16-17) homozygous for our modified CRISPR alleles (Fig. 2). We also used an antibody against horseradish peroxidase (anti-HRP, which recognizes a pan-neural epitope in the *Drosophila* central nervous system) to label all of the axons in the VNC and reveal the overall architecture of the axon scaffold.

**Figure 2.**
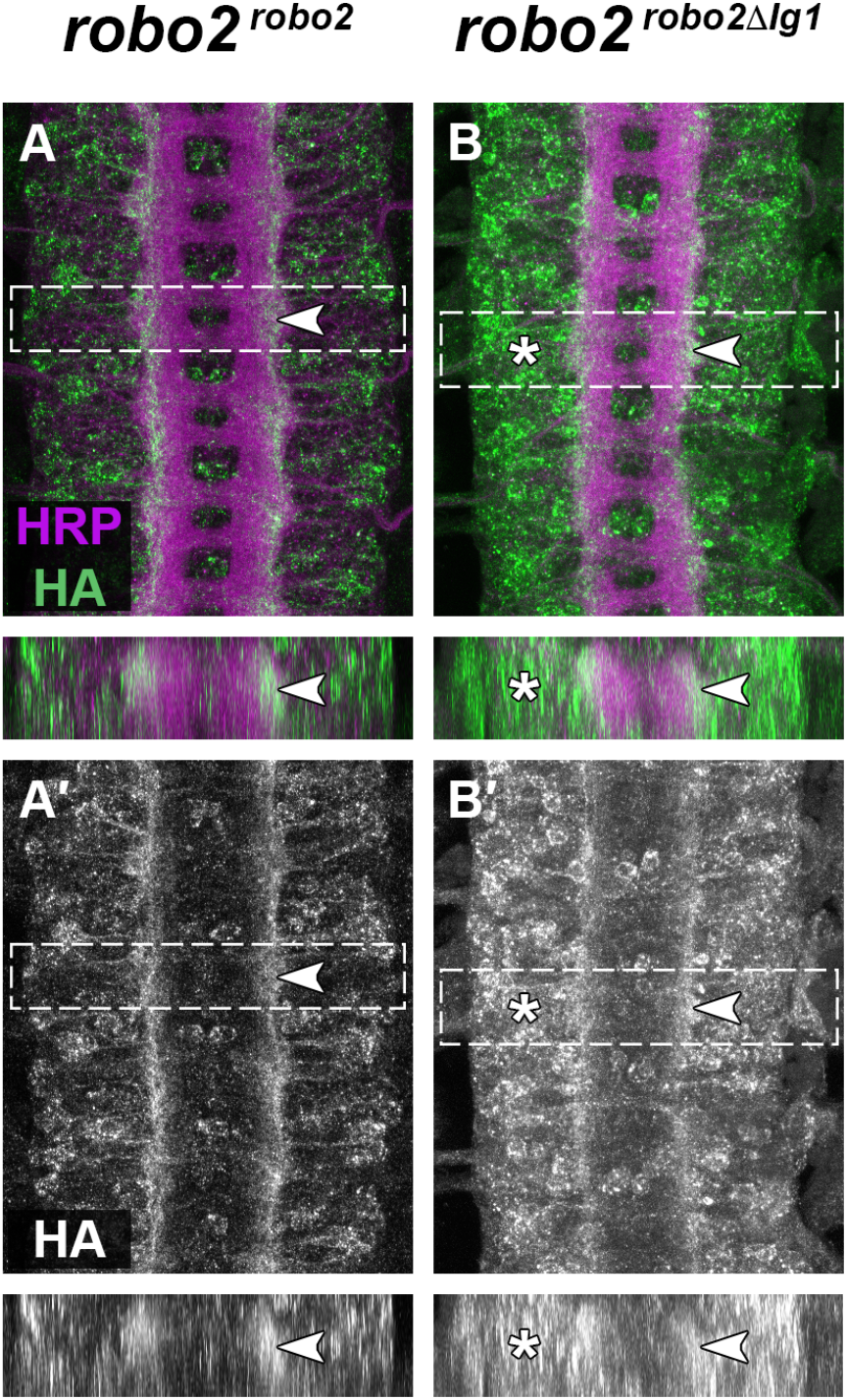
Expression of HA-tagged *robo2* alleles in the embryonic CNS. (A,B) Stage 16 *Drosophila* embryos stained with anti-HRP (magenta; labels all axons) and anti-HA (green) antibodies. (A′,B′) Anti-HA channels alone from the same embryos in (A,B). Lower panels show optical cross-sections of the regions outlined in dashed boxes. (A,A′) In *robo2*^*robo2*^ homozygous embryos, HA-tagged Robo2 protein reproduces Robo2’s endogenous expression pattern. At stage 16, Robo2 protein is primarily localized to longitudinal axons and restricted to the lateral-most region of the ventral nerve cord neuropile (arrowhead). (B,B′) In *robo2*^*robo2ΔIg1*^ homozygous embryos, HA-tagged Robo2ΔIg1 protein is detectable on lateral longitudinal axons (arrowhead) and also present at elevated levels on or in neuronal cell bodies within the cortex (asterisk).

Full-length Robo2 protein expressed from the *robo2*^*robo2*^ allele reproduces Robo2’s normal expression pattern in the ventral nerve cord of late-stage embryos: the protein is primarily localized to neuronal axons and restricted to the lateral-most longitudinal axon pathways in the neuropile (Fig. 2A) (Rajagopalan *et al*. 2000b; Simpson *et al*. 2000a). We are unable to directly compare our HA-tagged *robo2*^*robo2*^ CRISPR allele expression with endogenous Robo2 expression, as there is no monoclonal anti-Robo2 antibody (unlike for *Drosophila* Robo1 and Robo3), and the original polyclonal anti-Robo2 antibodies (Rajagopalan *et al*. 2000b; Simpson *et al*. 2000a) are no longer available. However, the expression pattern we observe closely matches previous descriptions of Robo2’s endogenous protein expression throughout embryogenesis, both in the VNC and other embryonic tissues (Rajagopalan *et al*. 2000b; Simpson *et al*. 2000a; Spitzweck *et al*. 2010). This result is also consistent with Spitzweck et al’s description of a similar knock-in *robo2*^*robo2*^ allele, and confirms that removing most of the introns from *robo2* and adding an N-terminal 4xHA tag does not interfere with the normal transcription or translation of *robo2*, or the stability, trafficking, or localization of the Robo2 protein (Spitzweck *et al*. 2010).

In homozygous *robo2*^*robo2ΔIg1*^ embryos, HA-tagged Robo2ΔIg1 protein was present on longitudinal axons and restricted to the lateral-most region of the neuropile, similar to full-length Robo2 (Fig. 2B, arrowhead). We also observed an increased degree of HA staining in neuronal cell bodies in the cortex surrounding the neuropile compared to *robo2*^*robo2*^ emrbyos (Fig. 2B, asterisk), suggesting that some portion of the Robo2ΔIg1 protein may not be trafficked correctly in embryonic neurons and instead retained at elevated levels in neuronal cell bodies. We also noted that the overall architecture of the axon scaffold appears affected in *robo2*^*robo2ΔIg1*^ homozygous embryos, with an overall decrease in the width of the scaffold along with irregularly shaped segmental neuromeres similar to *robo2* loss-of-function mutants, suggesting that replacing Robo2 with Robo2ΔIg1 may interfere with one or more aspects of neural development in *robo2*^*robo2ΔIg1*^ embryos.

### Robo2ΔIg1 cannot substitute for Robo2 to promote midline repulsion or lateral pathway formation

To examine specific axon guidance outcomes in *robo2*^*robo2*^ and *robo2*^*robo2ΔIg1*^ embryos, we used an anti-FasII antibody to label a subset of longitudinal axon pathways in the VNC. Robo2 is required for guidance of FasII-positive axons in the contexts of midline repulsion and longitudinal pathway formation: in *robo2* mutants, medial FasII-positive axons ectopically cross the midline (reflecting a lack of midline repulsion) and FasII-positive lateral axon pathways fail to form correctly (Rajagopalan *et al*. 2000a; Simpson *et al*. 2000b; Rajagopalan *et al*. 2000b; Simpson *et al*. 2000a). We quantified ectopic midline crossing and lateral pathway defects in *robo2*^*robo2*^ and *robo2*^*robo2ΔIg1*^ embryos stained with anti-FasII and anti-HRP, compared to *robo2* null mutants and heterozygous *robo2* control embryos (Fig. 3).

**Figure 3.**
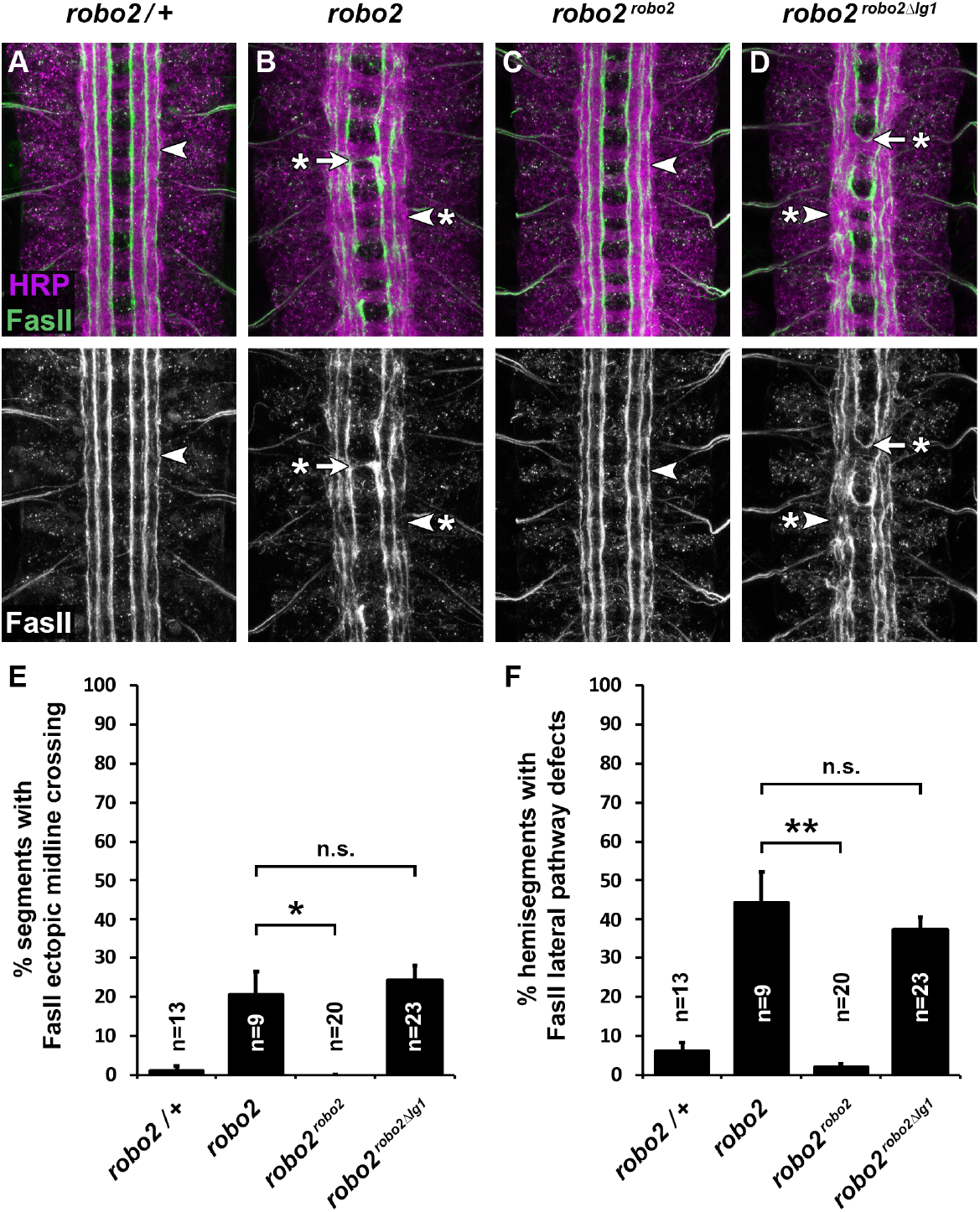
The Robo2 Ig1 domain is required for midline repulsion and lateral pathway formation. (A-D) Stage 16 *Drosophila* embryos stained with anti-HRP (magenta) and anti-FasII (green) antibodies. Lower images show anti-FasII channel alone from the same embryos. (A) In *robo2/+* heterozygous embryos, FasII-positive axons form three distinct longitudinal pathways on either side of the midline, one each in the medial, intermediate, and lateral zones of the neuropile. FasII-positive axons do not cross the midline in these embryos. Arrowhead points to the lateral FasII pathway. (B) In *robo2* loss of function mutants (*robo2*^*123*^*/robo2*^*135*^), FasII-positive axons cross the midline inappropriately in around 20% of segments (arrow with asterisk), and the lateral FasII pathway fails to form correctly in around 44% of hemisegments (arrowhead with asterisk). (C) In homozygous *robo2*^*robo2*^ embryos, midline repulsion and lateral pathway formation occurs normally. (D) Homozygous *robo2*^*robo2ΔIg1*^ embryos display ectopic midline crossing (arrow with asterisk) and lateral pathway defects (arrowhead with asterisk) equivalent to *robo2* mutants. (E,F) Quantification of ectopic midline crossing defects (E) and lateral pathway defects (F) in the genotypes shown in (A-D). Number of embryos scored for each genotype (n) is shown. Error bars indicate s.e.m. Percent defects for the two modified alleles were compared to *robo2* mutants by two-tailed Student’s t-test with a Bonferroni correction for multiple comparisons (*p<0.01; **p<0.001; n.s., not significant).

In heterozygous control *(robo2/+)* embryos, FasII-positive longitudinal pathways form correctly in three distinct zones within the neuropile of the embryonic VNC (medial, intermediate, and lateral), and FasII-positive axons do not cross the midline (Fig. 3A). In *robo2* amorphic mutant embryos *(robo2*^*123*^*/robo2*^*135*^), we observed ectopic midline crossing of FasII-positive medial axons in 20.6% of abdominal segments (segments A1-A7), and breaks in the lateral pathway and/or fusions between the lateral and intermediate pathways in 44.4% of abdominal hemisegments (left and right sides of segments A1-A7) (Fig. 3B). Neither of these defects are present in embryos homozygous for our *robo2*^*robo2*^ modified allele, indicating that expression of the HA-tagged full-length *robo2* cDNA in this allele can fully rescue *robo2-*dependent midline repulsion and longitudinal pathway formation (Fig. 3C). This result is consistent with a previous study by Spitzweck et al (2010), which reported that a *robo2*^*robo2*^ allele created via an ends-in knock-in approach could also fully rescue *robo2-*dependent axon guidance outcomes (Spitzweck *et al*. 2010).

In contrast, we observed ectopic midline crossing (24.2% of segments) and lateral pathway defects (37.6% of hemisegments) in *robo2*^*robo2ΔIg1*^ homozygous embryos at frequencies that were statistically indistinguishable from those in *robo2* amorphic mutants (p=0.42 and p=0.62 by t-test, respectively), suggesting that Robo2ΔIg1 is not able to substitute for full-length Robo2 in the contexts of midline repulsion or lateral pathway formation. The observation that lateral FasII-positive axon pathways are defective in *robo2*^*robo2ΔIg1*^ embryos, while Robo2-positive axons appear to be positioned correctly within the lateral zone, suggests that lateral positioning of FasII-positive and Robo2-positive lateral axons may occur independently.

## Discussion

Here we have described a CRISPR/Cas9-based gene replacement approach to characterize the functional importance of structural elements in the *Drosophila* Robo2 axon guidance receptor, and used this approach to show that the Slit-binding Ig1 domain of Robo2 is required for two distinct axon guidance roles of Robo2 during development of the *Drosophila* embryonic ventral nerve cord (midline repulsion and longitudinal pathway formation). We have also shown that the Ig1 domain contributes to the proper localization of Robo2 in embryonic neurons, suggesting a possible role for Robo2 Ig1 in protein trafficking to and/or retention in neuronal axons that is not conserved in *Drosophila* Robo1. The tools and approach we describe here will facilitate additional structure/function and gene replacement studies of *Drosophila robo2*.

### CRISPR gene replacement vs rescue transgene studies of *Drosophila* Robo receptors

We have previously used a transgene-based approach to characterize the functional importance of individual ectodomain elements in the *Drosophila* Robo1 protein (Brown *et al*. 2015; Reichert *et al*. 2016; Brown *et al*. 2018; Brown and Evans 2020). This approach relies on a rescue transgene carrying a small region of genomic DNA (∼4.5 kb) containing regulatory sequences sufficient to recapitulate the full expression pattern of *robo1*. Equivalent regulatory sequences have not been identified for *robo2* or *robo3*, so we could not use a similar rescue transgene approach for structure/function studies of Robo2. We have previously used a bacterial artificial chromosome (BAC) rescue approach employing a large (83.9 kb) *robo2-*containing BAC to examine the role of Robo2’s Ig2 domain in promoting midline crossing (Evans *et al*. 2015).

The CRISPR/Cas9-based strategy described here has a number of advantages over the above approaches, including: 1) identifying/isolating regulatory sequences is not required, as endogenous regulatory sequences are used instead; 2) the genetics of introducing markers and/or other mutations into the modified background is simplified, as there is no need to track an inactivating mutation plus a separate rescue transgene; and 3) the laborious recombineering and difficult transgenesis with very large BAC DNA fragments can be avoided. This CRISPR/Cas9 gene replacement approach could also be used to replace *robo2* with other coding sequences, including its paralogs from *Drosophila* (*robo1* and *robo3*), orthologs from other species, or chimeric/variant receptors, which would facilitate further structure/function or comparative/evo-devo studies. For example, we have used an equivalent approach to replace *Drosophila* Robo3 with its *Tribolium* ortholog Robo2/3 in order to compare their axon guidance activities (Evans 2017).

### Differential requirement for Ig1 in axonal localization of Robo1 and Robo2

We have previously reported that the Ig1 domain of *Drosophila* Robo1 is not required for proper expression or axonal localization of the Robo1 protein in the embryonic ventral nerve cord (Brown *et al*. 2015). The results presented here indicate that this is not true for *Drosophila* Robo2; instead, deleting Ig1 from Robo2 does appear to alter its subcellular localization in embryonic neurons. We see a similar effect on protein localization when the Ig1 domain is deleted from *Drosophila* Robo3 (Abigail Carranza and T.A.E., unpublished), suggesting that the Ig1 domains in Robo2 and Robo3 play a role in protein localization that is not shared by the Ig1 domain in Robo1. Importantly, deleting the Ig1 domain from Robo2 does not appear to affect its translation or protein stability, as Robo2 and Robo2ΔIg1 proteins are expressed at equivalent levels and detectable at the cell surface in cultured *Drosophila* S2R+ cells (Evans *et al*. 2015), and both proteins are detectable both on neuronal axons and in neuronal cell bodies *in vivo*, with the main difference being the relative levels in/on axons vs cell bodies (Fig. 2).

We have reported that deleting the Ig3 or Fn1 domains from Robo1 also resulted in elevated cell body expression of Robo1 (Reichert *et al*. 2016; Brown *et al*. 2018), but this effect (increased punctate staining in neuronal cell bodies for both Robo1ΔIg3 and Robo1ΔFn1) appears qualitatively distinct from what we observe with Robo2ΔIg1, where the increased cell body signal appears more membrane-localized (see Fig. 2B, where circular staining patterns presumably reflecting outlines of individual cell bodies can be seen). Whether this reflects differential roles for Robo2 Ig1 versus Robo1 Ig3/Fn1 in protein localization, or instead reflects underlying differences in the normal expression of Robo1 (which normally does not appear to reach the membrane in neuronal cell bodies) versus Robo2 (which is normally detectable at low levels on cell body membranes; see Fig. 2A), is unclear.

### Slit-dependent vs Ig1-dependent roles of Robo2

Although it is clear that Robo2’s normal roles in midline repulsion and lateral pathway formation are deficient in *robo2*^*robo2ΔIg1*^ embryos, we cannot distinguish between a direct requirement for Ig1 in each of these roles versus a secondary effect of altering Robo2’s subcellular distribution when Ig1 is deleted. In other words, perhaps Robo2ΔIg1 would be able to rescue some or all of these roles if it were primarily localized to axons similar to full-length Robo2.

We also note that while deleting the Ig1 domain from Robo2 does strongly or completely abrogate Slit binding (Evans *et al*. 2015), we cannot formally rule out the possibility that Ig1 may have other, Slit-independent activities that would also be disrupted by deleting the entire Ig1 domain. Slit binding in *Drosophila* and human Robo receptors can be strongly disrupted *in vitro* through targeted point mutations in Ig1 (Morlot *et al*. 2007; Fukuhara *et al*. 2008); a similar strategy might allow targeted disruption of Slit binding without altering other putative functions of Robo2 Ig1, which may in turn allow a more precise dissection of Slit-dependent vs Slit-independent roles of Ig1 *in vivo*. The CRISPR/Cas9-based gene replacement approach described here could be used to engineer a *robo2* locus expressing a cDNA carrying one or more targeted point mutations in Ig1 in order to test this possibility, and may also help disentangle the functional importance of Slit-binding versus axonal-localization activities of Robo2 Ig1.

### Does Robo2 guide FasII-positive longitudinal axons non-autonomously?

When *robo2’s* roles in embryonic axon guidance were first described two decades ago, the initial models posited that it acted as a cell-autonomous Slit receptor to signal midline repulsion and to guide longitudinal axons to lateral pathways (Rajagopalan *et al*. 2000a; Simpson *et al*. 2000b; Rajagopalan *et al*. 2000b; Simpson *et al*. 2000a). Subsequent studies of Robo2’s unique pro-crossing function revealed that Robo2 acts non-autonomously to guide commissural axons across the midline (Evans *et al*. 2015). We note that although FasII-positive lateral axons are misguided in *robo2*^*robo2ΔIg1*^ embryos, HA-positive axons in these embryos (that is, the axons that normally express Robo2) are still tightly restricted to the lateral region of the neuropile, and are not apparently misguided into the intermediate or medial zones. We have observed that HA-positive axons similarly remain restricted to the lateral zone in *robo2*^*robo1*^ embryos, in which the *robo2* coding sequence has been replaced by *robo1*, even though these embryos also display lateral FasII pathway defects similar to *robo2* null mutants (Spitzweck *et al*. 2010) (T.A.E., unpublished). These observations suggest that the FasII-positive axons that are misguided in *robo2*^*robo1*^ and *robo2*^*robo2ΔIg1*^ embryos (and *robo2* null mutants) may not be the same as the lateral axons that normally express Robo2. In other words, Robo2 may also act non-autonomously to regulate lateral position of FasII-positive axon pathways. Distinguishing between these possibilities will require examining both FasII and HA expression in the same embryos (which technical limitations have thus far prevented us from doing), and/or generating additional markers to label Robo2-expressing lateral axons independently of Robo2 expression in order to examine their lateral positions in wild-type, *robo2* mutant, and *robo2* gene replacement embryos. These results also demonstrate that the Ig1 domain of Robo2 is not required for guidance of Robo2-expressing longitudinal axons to lateral pathways, which indicates that the lateral axons that normally express Robo2 may not require its activity to form and/or join lateral pathways, or that Robo2 directs the medial-lateral positioning of these axons through an Ig1-independent mechanism.

## Materials and methods

### Molecular biology

#### Construction of robo2 donor plasmids

The *robo2*^*robo2*^ donor construct was assembled from four PCR fragments via Gibson assembly (New England Biolabs #E2611). The four fragments were derived from pBluescript (plasmid backbone), the wild type *robo2* genomic locus (5’ and 3’ homology regions), and an HA-tagged *robo2* cDNA plasmid (4xHA epitope tag and *robo2* coding region). The *robo2* coding sequence in the *robo2*^*robo2*^ donor construct is flanked by NheI and NotI restriction sites. To make the *robo2*^*robo2ΔIg1*^ donor, the full-length *robo2* coding sequence was excised with NheI-NotI and replaced with the *robo2ΔIg1* coding sequence. Donor plasmids contain engineered mutations in PAM and/or protospacer sequences to prevent cleavage by Cas9. Modified *robo2* HDR alleles include the following amino acid residues after the N-terminal 4xHA epitope tag, relative to Genbank reference sequence AAF51375: *robo2*^*robo2*^ (E89-V1463), *robo2*^*robo2ΔIg1*^ (E89-N90/L187-V1463). The entire donor regions including coding sequences and *robo2* flanking regions were sequenced prior to injection.

#### Construction of robo2 gRNA plasmid

*robo2* gRNA sequences were cloned into the tandem expression vector pCFD4 (Port *et al*. 2014) via PCR followed by Gibson assembly using the PCR product and BbsI-digested pCFD4 backbone. For gRNA 2, an additional G nucleotide was added to the 5’end of the gRNA target sequence to facilitate transcription from the U6-1 promoter.

### Genetics

#### Drosophila strains

The following *Drosophila* strains, transgenes, and mutant alleles were used: *robo2*^*robo2*^ and *robo2*^*robo2ΔIg1*^ (this study), *robo2*^*123*^ and *robo2*^*135*^ (Simpson *et al*. 2000b), *w*^*1118*^; *sna*^*Sco*^*/CyO,P{en1}wg*^*en11*^ *(Sco/CyOwg), y*^*1*^ *M{w[+mC]=nos-Cas9*.*P}ZH-2A w* (nos-Cas9*.*P)* (Port *et al*. 2014). All crosses were carried out at 25°C.

#### Generation and recovery of CRISPR modified alleles

The *robo2* gRNA plasmid was co-injected with the *robo2*^*robo2*^ or *robo2*^*robo2ΔIg1*^ homologous donor plasmids into *nos-Cas9*.*P* embryos (Bloomington *Drosophila* Stock Center stock #54591) (Port *et al*. 2014) by BestGene Inc (Chino Hills, CA). Injected individuals (G0) were crossed as adults to *Sco/CyOwg*. Founders (G0 flies producing F1 progeny carrying modified *robo2* alleles) were identified by testing two pools of three F1 females per G0 cross by genomic PCR with primers 458 and 325 (for *robo2*^*robo2*^) or 458 and 327 (for *robo2*^*robo2ΔIg1*^), which produce 1.5 kb products only when the respective HDR alleles are present. From each identified founder, 5-10 F1 males were then crossed individually to *Sco/CyOwg* virgin females. After three days, the F1 males were removed from the crosses and tested by PCR with the same set of primers to determine if they carried the modified allele. F2 flies from positive F1 crosses were used to generate balanced stocks, and the modified alleles were fully sequenced by amplifying the entire modified locus (approx. 6 kb) from genomic DNA using primers 458 and 545 or 458 and 594, then sequencing the PCR product after cloning via CloneJET PCR cloning kit (Thermo Scientific).

### Immunofluorescence and imaging

*Drosophila* embryo collection, fixation and antibody staining were carried out as previously described (Patel 1994). The following antibodies were used: mouse anti-Fasciclin II (Developmental Studies Hybridoma Bank [DSHB] #1D4, 1:100), mouse anti-βgal (DSHB #40-1a, 1:150), mouse anti-HA (BioLegend #901503, 1:1000), FITC-conjugated goat anti-HRP (Jackson Immunoresearch #123-095-021, 1:100), Alexa 488-conjugated goat Anti-HRP (Jackson Immunoresearch #123-545-021, 1:500), Cy3-conjugated goat anti-mouse (Jackson #115-165-003, 1:1000). Embryos were genotyped using balancer chromosomes carrying lacZ markers. Ventral nerve cords from embryos of the desired genotype and developmental stage were dissected and mounted in 70% glycerol/PBS. Fluorescent confocal stacks were collected using a Leica SP5 confocal microscope and processed by Fiji/ImageJ (Schindelin *et al*. 2012) and Adobe Photoshop software.

## Acknowledgments

Stocks obtained from the Bloomington Drosophila Stock Center [National Institutes of Health (NIH) grant P40 OD-018537) were used in this study. Monoclonal antibodies were obtained from the Developmental Studies Hybridoma Bank, created by the Eunice Kennedy Shriver National Institute of Child Health and Human Development of the NIH and maintained at The Department of Biology, University of Iowa, Iowa City, IA 52242. This work was supported by NIH grant R15 NS-098406 (T.A.E.) and by funds from the Arkansas Biosciences Institute (ABI).

